# Inhibition of the TPL2-MKK1/2-ERK1/2 pathway has cytostatic effect on B-Cell Lymphoma

**DOI:** 10.1101/2022.06.13.495940

**Authors:** Mariana Asslan, Guy Martel, Simon Rousseau

**Affiliations:** The Meakins-Christie Laboratories at the Research Institute of the McGill University Health Centre, McGill University, Montreal, Quebec, Canada; Department of Medicine, McGill University, Montreal, Quebec, Canada

**Keywords:** Blood cancer, Hematologic malignancies, lymphocytes

## Abstract

Diffuse Large B-Cell Lymphoma (DLBCL) are the most common form of non-Hodgkin lymphoma. Their molecular origin is heterogeneous and therefore treatments aimed at DLBCL must be adapted in function of the underlying molecular mechanisms driving cellular transformation. Constitutive activation of the protein kinases ERK1/2 is a hallmark of many B-cell malignancies. ERK1/2 activation which can occur downstream of the classical MAPK cascade via RAF or, in response to TLR stimulation, via the Tumor Promoting Locus 2 (TPL2) protein kinase. This pathway also relays signals from the MYD88 oncogenic mutant L265P, frequently found in hematologic malignancies. We report here that TPL2 participate to ERK1/2 activation downstream of BCR in a DLBCL cell line (OCI-Ly2). Moreover, we showed that a ERK2[Y316F] mutant increased *c-Myc*-luciferase reporter expression. We then investigated the impact of ERK1/2 inhibition on the proliferation of OCI-Ly2 cells. We found that blocking ERK1/2 MAPK signaling cascade using either MKK1/2 inhibitors (PD184352 and MEK162) or TPL2 inhibitor (Compound 1) was mainly cytostatic. Finally, we showed that while TPL2-MKK1/2 inhibition leads to cytostatic effect, Compound 1 has cytocidal effect at high concentrations, that is mediated via additional targets. Taken together, this study demonstrates the involvement of TPL2 in oncogenic signaling of DLBCL and supports the idea that combination therapy targeting multiple molecular pathways linked to cellular transformation is a superior avenue for future therapies.

## 2 Introduction

Constitutive activation of the protein kinases ERK1/ERK2 is a hallmark of many B-cell malignancies (Platanias, 2003a). Activation of ERK1/ERK2 can occur downstream of the Tumor Promoting Locus 2 (TPL2) protein kinase following activation of the Toll-like-receptor signaling pathway (Dumitru et al., 2000), the most frequently mutated signaling pathway in lymphoid neoplasms (Rousseau and Martel, 2016). This makes TPL2 an attractive target to treat hematologic malignancies.

Diffuse Large B-Cell Lymphoma (DLBCL) are the most common form of non-Hodgkin lymphoma. Their molecular origin is heterogeneous, and future therapies are aimed at targeting the molecular defects responsible for oncogenic progression in specific subsets. Molecular heterogeneity means that tumor cells employ different pathways to escape cell death signals and proliferate. To explore the relationship between molecular defects and drug-responsiveness, we delineated molecular pathways that are affected in a DLBCL cell line, OCI-Ly2 cells (Tweeddale et al., 1987). The cells were isolated from a 50-year-old male at relapse, with TP53 deletion that were MYC negative. They express low levels of the anti-apoptotic MCL1 protein but high levels of BCL-2, which correlates with their responsiveness to the BCL-2 inhibitors ABT-199 (Klanova et al., 2016). This pathway provides the necessary signals to escape cell death. We hypothesize that proliferative signals may be mediated by disturbances in the ERK1/ERK2 MAPK cascade. This would suggest that protein kinase inhibitors targeting this cascade may show anti-proliferative effect on OCI-Ly2 cells.

Activation of ERK1/ERK2 in hematologic malignancies can occur downstream of the classical RAS-RAF-MKK1/MKK2 pathway or alternatively downstream of the Tumor Promoting Locus 2 (TPL2) protein kinase (Rousseau and Martel, 2016). This pathway can relay signals to the ERK1/ERK2 MAPK from Toll-like Receptors and the MYD88 oncogenic mutant L265P (Dumitru et al., 2000; Beinke et al., 2004; Rousseau and Martel, 2016). ERK1/2 activation leads to the up regulation of several downstream effectors including c-Myc (Grandori et al., 2000). The c-Myc protein belongs to a family of transcriptional regulators and plays a critical role in controlling varied cellular processes such as growth and proliferation. Previously, we identified ERK2 mutations in hematologic malignancies, however, it is still not clear if these mutations can act themselves as tumorigenesis via the activation of c-Myc and other downstream effectors. Moreover, ERK1/2 can be activated by multiple upstream signaling pathways, in addition to the Toll-like Receptors’ pathway, such as B-cell Receptor (BCR) signaling. Previous studies indicated that ERK1/2 are involved in BCR-mediated expression of c-Myc (Moyo et al., 2017). However, it is not known whether the activation of BCR leads to ERK1/2 phosphorylation via TPL2 and the classical MAPK pathway.

In this report, we studied the effectiveness of TPL2 and the classical MAPK pathway inhibition on cell viability in non-Hodgkin lymphoma.

## 3 Methods

### 3.1. Cell culture

Immortalized human Diffuse Large B-Cell Lymphoma OCI-Ly2 cells were graciously provided by Prof. Minden (University of Toronto). RAMOS cells (Burkitts B-cell lymphoma) and HEK293 cells (epithelial kidney cells) were purchased from ATCC (Rockville, MD, USA). OCI-Ly2, RAMOS, and HEK293 cells were cultured in IMDM, RPMI 1640, and DMEM medium, respectively. Cells were supplemented with 10% fetal bovine serum and 1% penicillin/streptomycin. All cells were maintained at 37°C in 5 % CO2, 100% humidity. The medium was changed every 48–72 h until cells were treated as described.

### 3.2. c-Myc luciferase assay

HEK293 cells were transfected with 200 ng of pGL4.28-c-Myc and 800ng of pCDNA3.1-ERK2[D162N], pCDNA3.1-ERK2[D291G], pCDNA3.1-ERK2[Y316F] or the empty vector. Cells were lysed with Promega’s reporter buffer and subjected to luminescence analysis.

### 3.3. ERK1/2 immunoblotting

OCI-Ly2 were seeded in a 12-wells plate at 2 × 10^6^ cells/mL in IMDM supplemented with 0.5% FBS. Cells were grown for 24h and pre-treated for 1h with vehicle (DMSO), 2µM Compound 1 (C1) or 1.25µM MEK162. Cells were left untreated or stimulated with 5µg/mL anti-IgGMA for 10 min. Cells were lysed, 30µg of lysates was subjected to SDS-PAGE. Immunoblotting was performed with an antibody that recognizes ERK1/ERK2 phosphorylated at Thr202/Tyr204 or an antibody that recognizes all forms of ERK1/ERK2. Quantitative analysis of the signal intensity, obtained with an antibody recognizing only the phosphorylated forms of ERK1/ERK2 normalized to the signal obtained with antibody that recognizes all forms of ERK1/ERK2, was performed using LiCor infrared Odyssey imaging system and expressed as fold induction (Phospho-ERK1/2 average).

### 3.4. Trypan blue exclusion assay

Cells were seeded in 12-well plates at a concentration of 250,000 cells/mL. Cells were treated with increasing concentrations of PD184352. The plates were incubated at 37 °C in a humidified atmosphere containing 5 % CO2. After 72 h of incubation, cells centrifuged and resuspended in PBS. Then trypan blue was added (1:1) and viable cells were counted under microscope using hemocytometer.

### 3.5. Flow cytometry analysis of cell death by propidium iodide (PI) staining

OCI-Ly2 and Ramos cells were seeded at 250,000 cells/ml and treated with PD184352 at IC50 (4 and 3 µM respectively) for 72 h. As a positive control, cells were treated with 1.6mM H2O2 for 24 h. Prior to analysis by flow cytometry (BD LSRFortessa X-20), cells incubated at dark with PI staining for 5 mins. A total of 10,000 events were analyzed. Data were analyzed using FlowJo software.

### 3.6. MTT cell proliferation assay

OCI-Ly2 and Ramos cells were seeded in 96-well plates at 50,000 cells per well and incubated with increasing concentration of Compound 1 (C1) for 96 h. In addition, OCI-Ly2 cells were treated with: 625nM of Doxorubicin, 625nM of Vincristine, 10µM of Cyclophosphamide or 1µg/mL of Rituximab. Cell number was assessed with Vybrant® MTT Cell Proliferation Assay Kit, according to the manufacturer protocol (Thermofisher Scientific, Mississauga, Canada).

## 4 Results

### 4.1. ERK2[Y316F] increases *c-Myc* luciferase activity

We have previously shown that the most common mutation found in hematologic malignancies, MYD88[L265P] leads to ERK1/2 activation via TPL2 in a heterologous expression system (Rousseau and Martel, 2016). Interestingly, we had identified ERK2 mutations in hematologic malignancies that did not co-exist with MYD88 or other TLR-signaling components mutations suggesting that they may act themselves as drivers of disease. Since an important target of the ERK1/2 MAPK pathway is the oncogene MYC (Yasuda et al., 2008), we tested the capacity of ERK2[D162N], ERK2[D291G] and ERK2[Y316F] mutants to induce a c-Myc-luciferase reporter when over-expressed individually. We found that only ERK2[Y316F] increased *c-Myc*-luciferase reporter expression (**Fig. 1**). This provides additional support to the link between ERK1/2, MYC and B-cell tumorigenesis (Pulverer et al., 1994; Platanias, 2003b; Yasuda et al., 2008).

**Figure 1.**
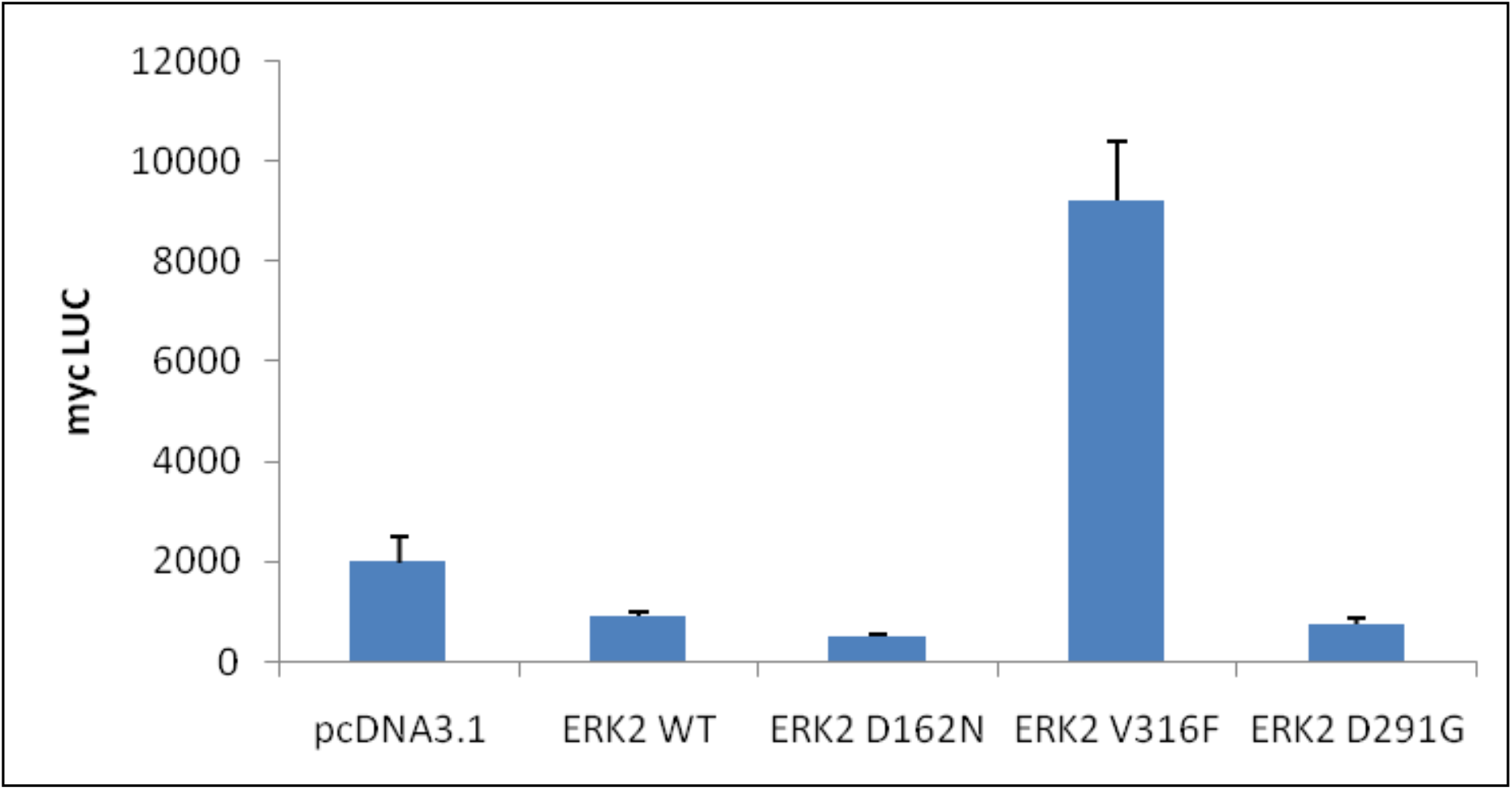
ERK2[Y316F] mutation increases c-Myc luciferase activity in HEK 293 cells. Cells were grown to confluence, lysed with Promega’s reporter buffer and subjected to luminescence analysis. All values are expressed as mean ± S.E.M. from four different experiments.

### 4.2. TPL2 and the classical MAPK pathway contribute to ERK1/ERK2 activation by BCR in B-cell lymphomas

B-cell Receptor (BCR)-mediated expression of MYC is regulated by ERK1/2 (Yasuda et al., 2008). However, it is not known whether physiological or abnormal activation of BCR, which activates the IKK complex, leads to ERK1/ERK2 phosphorylation via TPL2, which may contribute to B-cell malignancies. To test the involvement of TPL2, we exposed OCI-LY2 cells to a derivative of naphthyridine-3-carbonitrile, Compound 1, shown to inhibit TPL2 and prevent ERK1/ERK2 phosphorylation (Hall et al., 2007). Compound 1 did not decrease basal ERK1/ERK2 phosphorylation but decreased by more than half the anti-IgGMA-driven ERK1/ERK2 phosphorylation (**Table 1**). The FDA approved MKK1/2 inhibitor (MEK162) significantly reduced basal and anti-IgGMA-mediated ERK1/ERK2 phosphorylation. Therefore, TPL2 and the classical MAPK pathway contribute to ERK1/ERK2 activation by BCR in these cells.

**Table 1.** Impact of MEK162 and Compound 1 on the phosphorylation of ERK1/ERK2 MAPK in OCI-Ly2 cells.

| Cell line | Inhibitor | Treatment | Kinase targeted | Phospho-ERK1/2 average ± sem | P value (vs IgGMA) |
| --- | --- | --- | --- | --- | --- |
| OCI-Ly2 | vehicle | untreated | — | 1.0 ± 0.12 | |
| | C1 | untreated | TPL2 | 0.99 ± 0.11 | |
| | MEK162 | untreated | MKK1/MKK2 | 0.46 ± 0.17 | |
| | vehicle | IgGMA | — | 21.7 ± 3.22 | |
| | C1 | IgGMA | TPL2 | 9.67 ± 0.71 | p < 0.01 |
| | MEK162 | IgGMA | MKK1/MKK2 | 0.88 ± 0.29 | p < 0.001 |

OCI-Ly2 were grown for 24h, then pre-treated for 1h with vehicle (DMSO), 2 μM Compound 1 (C1) or 1.25 μM MEK162. Cells were left untreated or stimulated with 5 μg/mL anti-IgGMA for 10 min. Immunoblotting and quantitative analysis were performed. The signal intensity obtained with an antibody recognizing only the phosphorylated forms of ERK1/2 was normalized to the signal obtained with antibody that recognizes all forms of ERK1/2 using LiCor infrared Odyssey imaging system and expressed as fold induction (Phospho-ERK1/2 average).

### 4.3. TPL2 and MKK1/2 inhibition results in cytostatic effect on B-cells isolated from lymphomas

We next studied the impact of TPL2 and MKK1/2 inhibition on a B-cells derived from lymphomas. OCI-Ly2 cells were treated with MKK1/2 inhibitor PD184352 (CI-1040). Result showed that PD184352 reduced cell proliferation in a dose dependent manner (**Fig. 2A**). Similar results obtained with Burkitt lymphoma cell line Ramos (**Fig. 2B**). In both cell lines, OCI-Ly2 and Ramos, PD184352 at IC50 (4 and 3 µM respectively) produced cytostatic effects but not cytocidal (**Fig. 2C**). To test the involvement of TPL2 in the proliferation of OCI-Ly2 cells, they were exposed to either MEK162 or Compound 1, which efficiently decreased cellular proliferation (at IC50 5µM) and induced death at higher doses (10µM) (**Fig. 3A**). These concentrations are at least two-fold higher than the concentration required to achieve maximal TPL2 inhibition and prevent ERK1/ERK2 phosphorylation. The effect of Compound 1 does not appear to be restricted to OCI-Ly2 cells, as similar trend was also observed in the Burkitt lymphoma cell line Ramos (**Fig. 3B**). At the highest dose tested (10µM), Compound 1 was more effective than MEK162 in both cell lines. Moreover, Compound 1 was almost as efficacious as Rituximab, Doxorubicin and Vincristine in decreasing OCI-Ly2 cell numbers (**Fig. 3C**), not only preventing proliferation but leading to cell death. Taken together these results indicate that while TPL2-MKK1/2 inhibition leads to cytostatic effect, Compound 1 has cytocidal effect at high concentrations, suggesting that it acts via an additional target.

**Figure 2.**
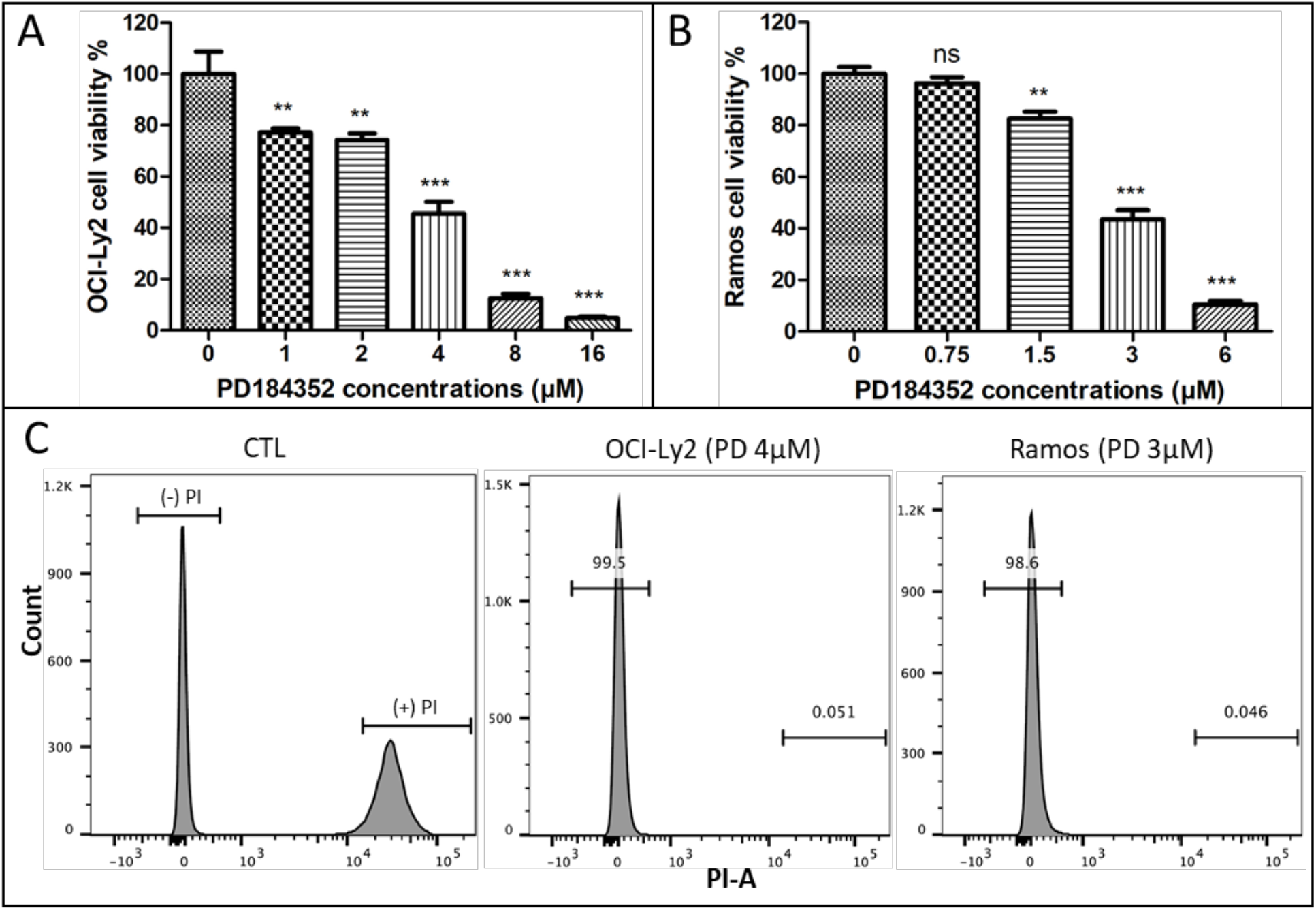
Impact of MKK1/2 inhibition on B-cell viability. **A**. OCI-ly2 cells or **B**. Ramos cells were seeded in 12-wells plate at 2.5× 10^5^ cells/mL and treated with increasing concentration of PD184352 for 72h. Each treatment was done in duplicates, and the experiments were done at least thrice. Cell viability was measured with Trypan Blue exclusion assay. Data presented as the mean ± S.E.M. (one-way ANOVA with Dunnett test ***, p < 0.001; **, p < 0.01; *, p < 0.05). **C**. Flow cytometry histograms showing propidium iodide (PI) staining after treatment with PD184352 at IC50. Either OCI-Ly2 cells (middle panel) or Ramos cells (right panel) were stained with PI after 72h of treatment as in **A** and **B**.

**Figure 3.**
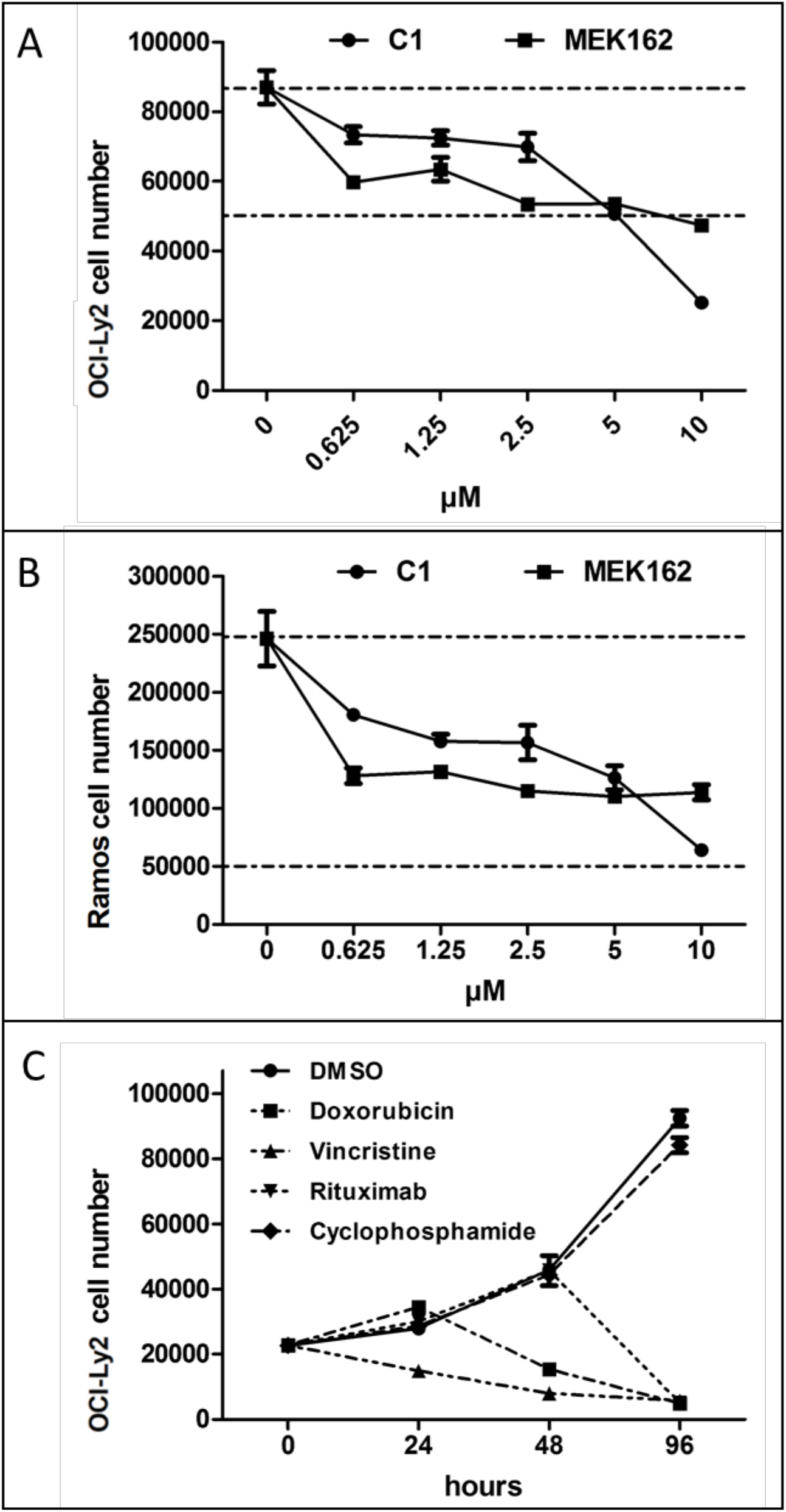
Impact of TPL2 inhibition on B-cell viability. **A**. OCI-ly2 cells were seeded in 96-well plates (50,000 cells per well) and incubated with increasing concentration of Compound 1 (C1) for 120h. Each treatment was done in quadruplicate wells, and the experiments were done at least thrice. Cell number was assayed at 72h with Vybrant® MTT Cell Proliferation Assay Kit. The results are expressed as the mean ± S.E.M. of a single representative experiment. Lower dashed lines represent the number of cells seeded while the top line represents the cell number at 120h in vehicle. **B**. Ramos cells were treated with C1 and cell number was measured after 48h of incubation as in **A. C**. OCI-Ly2 were incubated with: 625nM of Doxorubicin, 625nM of Vincristine, 10µM of Cyclophosphamide or 1µg/mL of Rituximab and cell number was assessed using MTT assay.

## 5 Discussion

In this report we confirmed that blocking ERK1/2 MAPK signaling cascade alone seems to be mainly cytostatic and its antitumor activity may not necessarily lead to tumor regression (Montagut and Settleman, 2009). Hence, additional therapeutic modality with antiapoptotic effectiveness may help to maximize the antitumor effectiveness of MAPK signaling inhibitors. It had been postulated that oncogenic kinase inhibitors efficacy may be boosted by the presence of BCL-2 inhibitors (Cragg et al., 2009). Furthermore, previous studies indicate that the therapeutic benefit of BCL-2 inhibitors alone such as ABT-199 and ABT-263 is limited due to acute, dose-dependent thrombocytopenia (Itchaki and Brown, 2016). Therefore, future studies should consider combination therapy between these two families or others that may yield higher and more durable responses and limiting the detrimental effects.

These results presented also show a contribution of TPL2 to ERK1/ERK2 activation following BCR activation in OCI-Ly2 cells. This finding has important implication for further investigation of compounds that aims to prevent ERK1/ERK2 activation following gain-of-function mutations downstream of BCR signaling. Both the classical RAF-activated MAPK cascade and the IKK-TPL2 pathways appears to contribute to the activation of ERK1/ERK2.

Finally, a surprising but very interesting result is the impact of Compound 1 at high concentrations on OCI-Ly2 cell numbers. At the highest dose tested (10µM), Compound 1 was almost as efficacious as Rituximab, Doxorubicin and Vincristine in decreasing OCI-Ly2 cell numbers, not only preventing proliferation but leading to cell killing. This effect is likely mediated by inhibition of other targets in addition to TPL2 and ERK1/ERK2. Whether this new target act in concert or independently to ERK1/ERK2 activation remains to be determined. This other target is likely another protein kinase, as Compound 1 is an ATP competitive inhibitor that came from a tyrosine kinase inhibitor collection (Green et al., 2007). Using the published profile of kinases affected by Compound 1 (Hall et al., 2007), the second most affect protein kinase is CAMKII. Interestingly, CAMKIIγ gamma stabilizes the c-Myc protein through its phosphorylation and that inhibition of CAMKII reduced tumor burden in T cell lymphomas (Gu et al., 2017). This raises the possibility that Compound 1 may decrease CAMKII activity at the highest concentrations used, reducing MYC stability. This could be sufficient to induce killing. More likely, Compound 1 may act via the combined action of the kinases it targets, preventing ERK1/2 induced c-Myc gene expression via TPL2 and MYC protein destabilization via CAMKII. BCR-mediated expression of *c-myc* is regulated via ELK-1 phosphorylation carried out by ERK1/2 (Yasuda et al., 2008). Further supporting the link between ERK1/2 and MYC in B-cell tumorigenesis, we report the first experimental assay demonstrating that ERK2[Y316F] has biological activity, leading to increased expression of a c-Myc luciferase reporter. Moreover, ERK1/2 can phosphorylate the same residue as CAMKII (Ser62) on MYC (Pulverer et al., 1994), raising the possibility that inhibition of one kinase can be compensated by the other, providing another path by which Compound 1 could be acting to induce cell killing.

As Compound 1 was develop for its selectivity towards TPL2, it may be interesting to revisit the original series of compounds and those of the related quinoline-3-carbonitriles series for their potency at killing DLBCL cells (Green et al., 2007; Hall et al., 2007). These other compounds may have greater potency towards lymphoma cell killing and enable the development of novel anti-lymphoma drugs that target lymphoma unresponsive to ERK1/2 MAPK cascade inhibition. Adding novel chemical families to the arsenal of drugs that can impact lymphomas will be important to target the molecular heterogeneity of tumor cells and provide patient specific therapies.

## Funding Statement

This work was supported by Canadian Institute of Health Research (MOP#123496) and the Fonds de Recherche du Québec -Santé (FRQ-S salary award to S.R.). The Meakins-Christie Laboratories – RI-MUHC, are supported by a Centre grant from FRQ-S.

## Notes

### Competing Interest Statement

The authors have declared no competing interest.

## References

Beinke, S., Robinson, M., Hugunin, M., and Ley, S. (2004). Lipopolysaccharide activation of the TPL-2/MEK/extracellular signal-regulated kinase mitogen-activated protein kinase cascade is regulated by IkappaB kinase-induced proteolysis of NF-kappaB1 p105. Molecular and cellular biology 24, 9658–67. doi: 10.1128/mcb.24.21.9658-9667.2004.

Cragg, M. S., Harris, C., Strasser, A., and Scott, C. L. (2009). Unleashing the power of inhibitors of oncogenic kinases through BH3 mimetics. Nature Reviews Cancer 9, 321–326. doi: 10.1038/nrc2615.

Dumitru, C., Ceci, J., Tsatsanis, C., Kontoyiannis, D., Stamatakis, K., Lin, J., et al. (2000). TNF-alpha induction by LPS is regulated posttranscriptionally via a Tpl2/ERK-dependent pathway. Cell 103, 1071–83.

Grandori, C., Cowley, S. M., James, L. P., and Eisenman, R. N. (2000). The Myc/Max/Mad Network and the Transcriptional Control of Cell Behavior. Annual Review of Cell and Developmental Biology 16, 653–699. doi: 10.1146/annurev.cellbio.16.1.653.

Green, N., Hu, Y., Janz, K., Li, H.-Q., Kaila, N., Guler, S., et al. (2007). Inhibitors of tumor progression loci-2 (Tpl2) kinase and tumor necrosis factor alpha (TNF-alpha) production: selectivity and in vivo antiinflammatory activity of novel 8-substituted-4-anilino-6-aminoquinoline-3-carbonitriles. Journal of medicinal chemistry 50, 4728–45.

Gu, Y., Zhang, J., Ma, X., Kim, B., Wang, H., Li, J., et al. (2017). Stabilization of the c-Myc Protein by CAMKIIγ Promotes T Cell Lymphoma. Cancer Cell 32, 115–128.e7. doi: 10.1016/j.ccell.2017.06.001.

Hall, J., Kurdi, Y., Hsu, S., Cuozzo, J., Liu, J., Telliez, J.-B. B., et al. (2007). Pharmacologic inhibition of tpl2 blocks inflammatory responses in primary human monocytes, synoviocytes, and blood. The Journal of biological chemistry 282, 33295–304. doi: 10.1074/jbc.M703694200.

Itchaki, G., and Brown, J. R. (2016). The potential of venetoclax (ABT-199) in chronic lymphocytic leukemia. Therapeutic advances in hematology 7, 270–287. doi: 10.1177/2040620716655350.

Klanova, M., Andera, L., Brazina, J., Svadlenka, J., Benesova, S., Soukup, J., et al. (2016). Targeting of BCL2 Family Proteins with ABT-199 and Homoharringtonine Reveals BCL2-and MCL1-Dependent Subgroups of Diffuse Large B-Cell Lymphoma. Clinical cancer research : an official journal of the American Association for Cancer Research 22, 1138–49. doi: 10.1158/1078-0432.CCR-15-1191.

Montagut, C., and Settleman, J. (2009). Targeting the RAF-MEK-ERK pathway in cancer therapy. Cancer letters 283, 125–34. doi: 10.1016/j.canlet.2009.01.022.

Moyo, T. K., Wilson, C. S., Moore, D. J., and Eischen, C. M. (2017). Myc enhances B cell receptor signaling in precancerous B cells and confers resistance to Btk inhibition. Oncogene 36, 4653–4661. doi: 10.1038/onc.2017.95.

Platanias, L. C. (2003a). Map kinase signaling pathways and hematologic malignancies. Blood 101, 4667–79. doi: 10.1182/blood-2002-12-3647.

Platanias, L. C. (2003b). Map kinase signaling pathways and hematologic malignancies. Blood 101, 4667–79. doi: 10.1182/blood-2002-12-3647.

Pulverer, B., Fisher, C., Vousden, K., Littlewood, T., Evan, G., and Woodgett, J. (1994). Site-specific modulation of c-Myc cotransformation by residues phosphorylated in vivo. Oncogene 9, 59–70.

Rousseau, S., and Martel, G. (2016). Gain-of-Function Mutations in the Toll-Like Receptor Pathway: TPL2-Mediated ERK1/ERK2 MAPK Activation, a Path to Tumorigenesis in Lymphoid Neoplasms? Frontiers in cell and developmental biology 4, 50. doi: 10.3389/fcell.2016.00050.

Tweeddale, M., Lim, B., Jamal, N., Robinson, J., Zalcberg, J., Lockwood, G., et al. (1987). The presence of clonogenic cells in high-grade malignant lymphoma: a prognostic factor. Blood 69, 1307–14.

Yasuda, T., Sanjo, H., Pagès, G., Kawano, Y., Karasuyama, H., Pouysségur, J., et al. (2008). Erk kinases link pre-B cell receptor signaling to transcriptional events required for early B cell expansion. Immunity 28, 499–508. doi: 10.1016/j.immuni.2008.02.015.

